# Cyst size variability in invasive American *Artemia franciscana* Kellogg, 1906 (Branchiopoda: Anostraca) in Asia: a commercial approach

**DOI:** 10.1101/2022.05.07.491002

**Authors:** Xiao-Fang Wu, Chun-Yang Shen, Chun-Zheng Fu, Ning Yang, Wang Pei-Zheng, Amin Eimanifar, Alireza Asem

## Abstract

*Artemia* is the most common live food which is used in aquaculture worldwide. This study reports on biometrical variation of introduced American *Artemia franciscana* cyst from 24 non-native localities and two native habitats in Asia and USA, respectively. Results showed the largest diameter of untreated cyst, diameter of decapsulated cyst and thicker chorion ordinarily belong to invasive populations. Because of the small cysts, which have an effect on increasing quantity per unit weight and could be the cause of increased hatching efficiency, commercial productions of *A. franciscana* cyst from native sources should potentially be considered higher quality than productions from non-indigenous environments. Principal Component Analysis revealed that all cyst batches from San Francisco Bay were classified in one group and the most invasive populations could arrange in other separated group. Although, diameter of decapsulated cyst and chorion thickness showed a negative and significant correlation among invasive populations, there was no significant relationship within native populations. These observations contrast with biometrical patterns of parthenogenetic populations.

## INTRODUCTION

Different productions of brine shrimp *Artemia* including newly hatched nauplii and decapsulated cysts, have been widely used in fishery and aquaculture industry (Sorgeloos et al., 2001). From 1980 to 1994, the annual consumption of *Artemia* cyst has rapidly increased from 60 tons to 2000 tons (Bengtson et al., 1991; Triantaphyllidis et al., 1994). In the early of 21th century, aquaculture development programs raised the harvesting wet *Artemia* cyst nearly 9000 tons from Great Salt Lake, USA (Dhont & Sorgeloos, 2002) Van Stappen et al. (2020) have estimated that consumption of *Artemia* cyst in China was 1000 tons in 2016.

Since the 1950s, to support the developing aquaculture industry, the cyst of American *Artemia franciscana* have been exported overseas from Great Salt Lake and San Francisco Bay in the USA (Van Stappen, 2008; Eimanifar et al., 2014). Due to provide the live food demanded in fishery and regarding to the adaptation ability of *A. franciscana* in the extreme environmental conditions, it has been cultured in non-native habitats and man-made salterns which has caused the permanent colonization in numerous geographical regions across Eurasia, Mediterranean regions (Amat et al., 2005; Eimanifar et al., 2014, 2020; Scalone and Rabet, 2013; Saji et al., 2019; Shen et al., 2021; Asem et al., 2021) and Australia (Asem et al., 2018).

Aquaculture industry is the basic and main reason of dispersal of American *A. franciscana* in non-native habitats especially in Asia (Camara, 2020; Shen et al., 2021). Although genetic variation of invasive *A. franciscana* has been well studied (Scalone and Rabet, 2013; Eimanifar et al., 2014, 2020; Saji et al., 2019; Saad & Elsebaie, 2020; Asem et al., 2018, 2021), there is a lack of information on biometrical variation of cyst in new environments. Beside of several biological factors including nutrition and hatching percentage, the hatching efficiency (the number of nauplii which obtain with per gram of dry cysts) is an important parameter in evaluation of cyst quality and its marketing (Sorgeloos et al., 1978). Even if, physicochemical conditions can affect hatching percentage as well as hatching efficiency (Camargo et al., 2004; Sayg, 2004; Salma et al., 2012; El-Magsodi et al., 2016; Sharahi & Zarei, 2016), *Artemia* cyst size has an impact on number of eggs in per unit weight which is an impressive parameter on hatching efficiency regardless of hatching conditions (Asem et al., 2007, 2010).

The aim of this study was to investigate the variation of introduced American *A. franciscana* cysts (resting eggs) in Asia. First, we compared biometrical variability within and between native and exotic populations to understand alteration patterns of cyst size in non-indigenous environments. Second, correlation of cyst size was analyzed with decapsulated (yolk sac and embryo) and chorion (outer shell of egg) thickness to determine the relationships among biometrical characters. Gaining a better apprehension of biometric characterization of *Artemia* cyst, could supply insight into the reasons why different batches/populations with same hatching percentage, represent different hatching efficiency.

## MATERIALS AND METHODS

Biometrical characterizations of invasive American *A. franciscana* cyst were studied in 24 localities from Asia and two native habitats in USA (Great Salt Lake, UT and San Francisco Bay, CA) (table 1).

**Table 1.** Biometric mean (±SD) of invasive and native populations of *A. franciscana* cysts from Asia and USA. Same letters in each column show non-significant difference (n = 100, ANOVA, Tukey test, p > 0.05).

| | Location<sup>1</sup> | Abb. | Untreated cyst (μm) | Decapsulated cyst (μm) | Chorion thickness (μm) |
| --- | --- | --- | --- | --- | --- |
| Invasive populations | Tanggu, China | TTA | 251.30±14.95<sup>fg</sup> | 233.90±14.70<sup>fg</sup><sup>hi</sup> | 8.70 |
| | Nanpu, China | NAN | 265.90±15.67<sup>h</sup> | 245.90±14.45<sup>l</sup> | 10.00 |
| | Yuantong, China | YUA | 245.60±12.21<sup>cdefg</sup> | 229.85±14.19<sup>defgh</sup> | 7.88 |
| | Zhan hua, China | ZHS | 241.70±11.94<sup>bcd</sup> | 218.55±10.55<sup>ab</sup> | 11.58 |
| | Wuzhidui, China | WUZ | 238.85±10.91<sup>bc</sup> | 229.45±13.44<sup>defg</sup> | 4.70 |
| | Bohai Bay, China | BBA | 247.63±14.38<sup>defg</sup> | 226.60±10.91<sup>cde</sup> | 10.51 |
| | Xinhu, China | XIN | 252.05±13.65<sup>g</sup> | 237.10±10.57<sup>ij</sup> | 7.48 |
| | Yanhua, China | YANH | 252.15±11.64<sup>g</sup> | 232.53±9.93<sup>efghi</sup> | 9.81 |
| | Haixing, China | HAI | 245.85±12.43<sup>defg</sup> | 228.65±13.52<sup>cdef</sup> | 8.60 |
| | Wudi, China | WUD | 244.75±12.72<sup>cdef</sup> | 227.25±10.60<sup>cde</sup> | 8.75 |
| | Beidaba, China | BEID | 247.45±12.03<sup>defg</sup> | 228.35±10.54<sup>cdef</sup> | 9.55 |
| | Leguantai, China | LEG | 249.70±15.95<sup>efg</sup> | 232.00±13.08<sup>efghi</sup> | 8.85 |
| | Dongying, China | DOG | 249.40±12.82<sup>efg</sup> | 228.2±13.06<sup>cdef</sup> | 10.60 |
| | Sikou, China | SIK | 251.30±14.15<sup>fg</sup> | 235.55±12.73<sup>ghij</sup> | 7.88 |
| | Yangkou, China | YAG | 267.90±19.15<sup>h</sup> | 250.55±15.83<sup>ml</sup> | 8.67 |
| | Da Gang, China | DAG | 235.90±14.34<sup>ab</sup> | 225.55±13.87<sup>cd</sup> | 5.18 |
| | Eryan, China | ERY | 249.95±14.24<sup>fg</sup> | 228.2±11.09<sup>cdef</sup> | 10.88 |
| | Vinhchau, Vietnam | VCH | 242.90±11.06<sup>cde</sup> | 222.65±8.92<sup>abc</sup> | 10.13 |
| | Sri Lanka | SLA | 241.50±11.23<sup>abc</sup> | 222.35±7.96<sup>ab</sup> | 9.58 |
| | Kelambakkam, India | KEL | 242.50±11.00<sup>bcd</sup> | 235.90±11.51<sup>hij</sup> | 3.30 |
| | Tuticorin, India | TUT | 249.65±10.67<sup>efg</sup> | 225.20±10.27<sup>cd</sup> | 12.23 |
| | Nough catchment, Iran | NOG | 251.00±13.96<sup>fg</sup> | 255.40±15.66<sup>m</sup> | 2.20 |
| | Mahshahr port, Iran | MAH | 246.00±14.34<sup>defg</sup> | 240.60±14.24<sup>jk</sup> | 2.70 |
| | Garmat Ali, Iraq | GAA | 241.05±11.13<sup>bcd</sup> | 224.00±8.67<sup>bcd</sup> | 8.53 |
| Native populations | Great Salt Lake, USA<sup>2</sup> | GSL | 262.50±13.04<sup>h</sup> | 245.15±9.55<sup>l</sup> | 8.68 |
| | San Francisco Bay, USA<sup>2</sup> | SFB | 230.85±10.71<sup>a</sup> | 216.40±12.00<sup>a</sup> | 7.23 |
| | San Francisco Bay, USA<sup>3*</sup> | SFB<sub>VS1</sub> | 224.70±12.4 | 210.00±12.7 | 7.35 |
| | San Francisco Bay, USA<sup>3*</sup> | SFB<sub>VS2</sub> | 224.60±11.9 | 210.50±12.3 | 7.05 |
| | San Francisco Bay, USA<sup>3*</sup> | SFB<sub>VS3</sub> | 223.90±11.7 | 209.70±12.8 | 7.10 |
| | San Francisco Bay, USA<sup>3*</sup> | SFB<sub>VS4</sub> | 224.30±11.8 | 207.70±11.1 | 8.30 |
| | San Francisco Bay, USA<sup>3</sup> | SFB<sub>VS5</sub> | 228.70±12.3 | 212.10±11.3 | 8.30 |
| | Great Salt Lake, USA<sup>3*</sup> | GSL<sub>VS1</sub> | 252.50±13.0 | 241.60±13.2 | 5.45 |
| | Great Salt Lake, USA<sup>3*</sup> | GSL<sub>VS2</sub> | 244.20±16.1 | 234.80±16.0 | 4.7 |
| | Great Salt Lake, USA<sup>4*</sup> | GSL<sub>E1</sub> | 230.99±11.32 | 213.69±11.16 | 8.65 |
| | Great Salt Lake, USA<sup>4*</sup> | GSL<sub>E2</sub> | 231.92±11.80 | 216.32±10.71 | 7.8 |
| | Great Salt Lake, USA<sup>4*</sup> | GSL<sub>E3</sub> | 216.17±9.15 | 214.03±10.01 | 1.07 |
| | Great Salt Lake, USA<sup>4*</sup> | GSL<sub>E4</sub> | 232.71±11.18 | 217.58±11.93 | 7.56 |
| | Great Salt Lake, USA<sup>4*</sup> | GSL<sub>E5</sub> | 233.06±9.83 | 217.65±10.90 | 7.7 |
| | Great Salt Lake, USA<sup>4*</sup> | GSL<sub>E6</sub> | 218.30±13.12 | 210.81±10.44 | 3.74 |

Note: 1) further information of studied localities are available in Eimanifar et al. (2014). 2) This study, 3) Vanhaecke and Sorgeloos, 1980, 4) Eimanifar et al., 2015.
\* These results of Vanhaecke and Sorgeloos (1980) and Eimanifar et al. (2015) have not been used for one-way ANOVA analyses.

In each population, cysts were hydrated and decapsulated following Asem et al. 2007. Diameters of untreated (fully cyst) and decapsulated cysts were measured using a Motic BA210 microscope equipped with a MoticamX camera and software of Motic Images Plus 2.0v. Thickness of chorion layer was determined by: chorion thickness = (Mean diameter of untreated cyst **-** Mean diameter of decapsulated cyst)**/**”. This value is reported without standard deviations (Vanhaecke & Sorgeloos, 1980; Asem et al., 2010, 2014).

One-Way ANOVA (Tukey test, p > 0.05) was performed to analyze significant differences among biometrical characters. Three parameters (diameter of untreated cysts, diameter of decapsulated cyst and chorion thickness) were employed for the clustering of populations using Principal Component Analysis (PCA). The patterns of cyst variability were analyzed using linear regression and Pearson’s correlation between the biometrical parameters (p > 0.05). The computer program SPSS 22 was utilized for statistical analysis.

## RESULTS

### Cyst Biometry

Biometrical characters of *A. franciscana* cyst is summarized in table 1. According to the findings, the biggest fully cyst of *Artemia* belongs to invasive population of YAG (267.90±19.15 µm) from China that has significant differences with other remaining regions sample, excepting invasive population of NAN (265.90±15.67 µm) and a batch of native population from GSL (262.50±13.04 µm). Among invasive populations, DAG (235.90±14.34 µm) and WUZ (238.85±10.91 µm) yields the smallest size of untreated cyst from China. Generally native populations (GSL and SFB) represent small fully cyst.

Two invasive populations, NOG and YAG reveal the largest diameter of decapsulated cysts (255.40±15.66 and 250.55±15.83 µm, respectively). While among invasive populations, the smallest decapsulated cyst belongs to ZHS (218.55±10.55 µm), overall the smallest decapsulated cyst are observed in native population from SFB (from 207.7±11.1 to 216.40±12.00 µm).

The thinnest chorion layers are observed in one of native population of GSL batches (1.07 µm), followed by invasive populations of NOG (2.20 µm) and MAH (2.70 µm) from Iran. The thickest chorion layer belongs to TUT (12.23 µm) from India, followed by ZHS (11.58 µm) from China, respectively.

### Principal Component Analysis

Regarding to the PCA (Principal Component Analysis), the first and second components present 64.18% and 35.66% of variability and totally the both components involve with 99.84% of separation. In the first component, the means of untreated cyst (0.996) and the means of decapsulated cyst (0.921), have the main influences in the classification of populations, respectively. The chorion thickness (0.956) is an effective character in the second component. PCA shows two separated collections, a group containing all batches of native population from SFB and four batches of GSL (Group A), and another consists of 17 invasive populations (Group B). While 12 populations (five batches of native populations from GSL and seven batches of invasive populations) show a widely distribution on PCA plot without grouping (fig. 1).

**Fig. 1.**
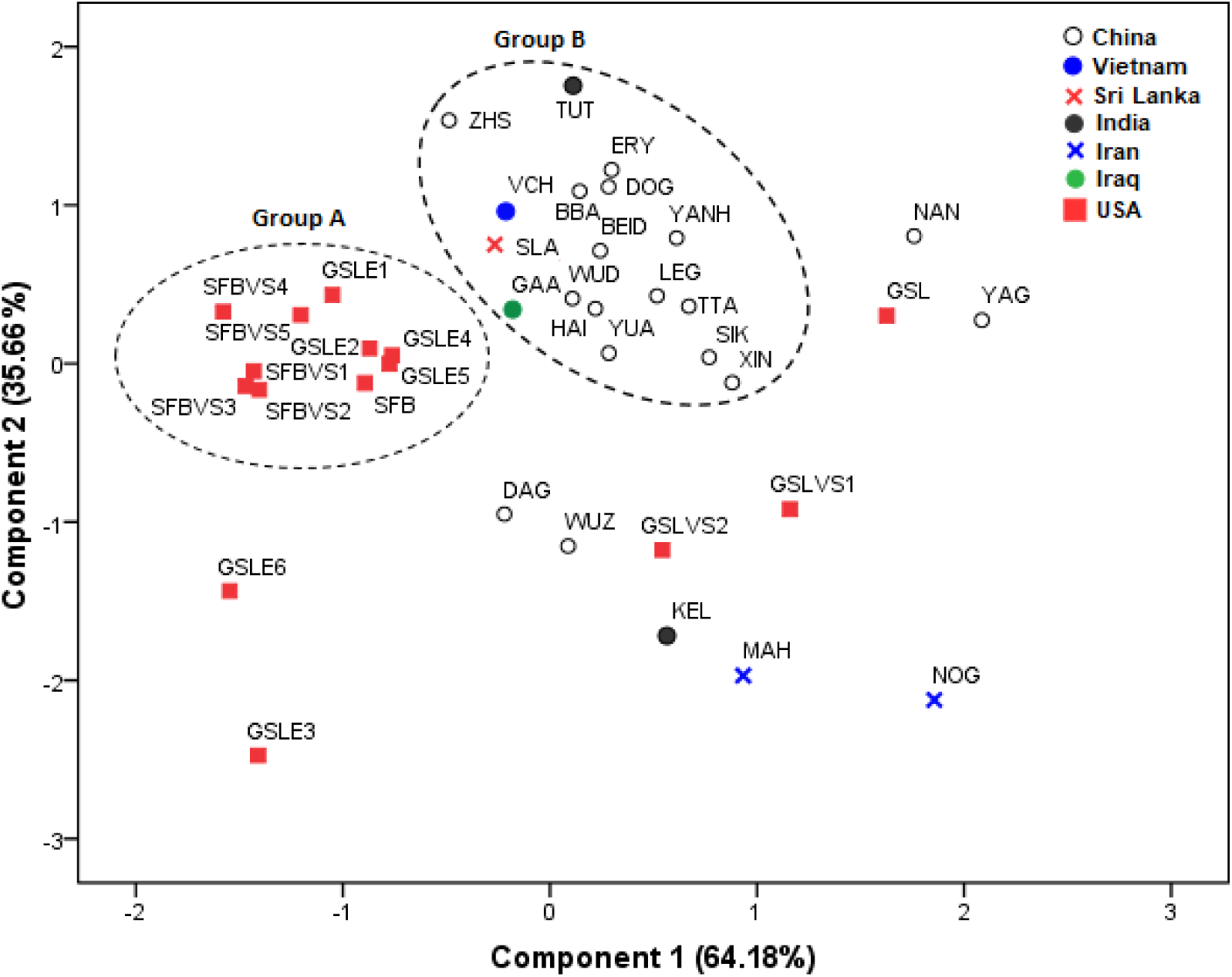
Scatterplot of Principal Component Analysis (PCA) based on three biometrical characters of cysts.

### Regression and Correlation

The equations of Linear Regression and Pearson’s correlations are showed in figure 2. The results confirm a positive and significant correlation between size of untreated and decapsulated cyst in both native and invasive populations (r = 0.941, Sig. = 0.0001 and r = 0.969, Sig. = 0.0001, respectively). On the other hand, there are no significant correlations between untreated cyst and chorion thickness. While correlation between decapsulated cyst and chorion thickness is non-significant in native populations (r = 0.041, Sig. = 0.883), invasive populations show negative and significant relationship (r = 0.521, Sig. = 0.009).

**Fig. 2.**
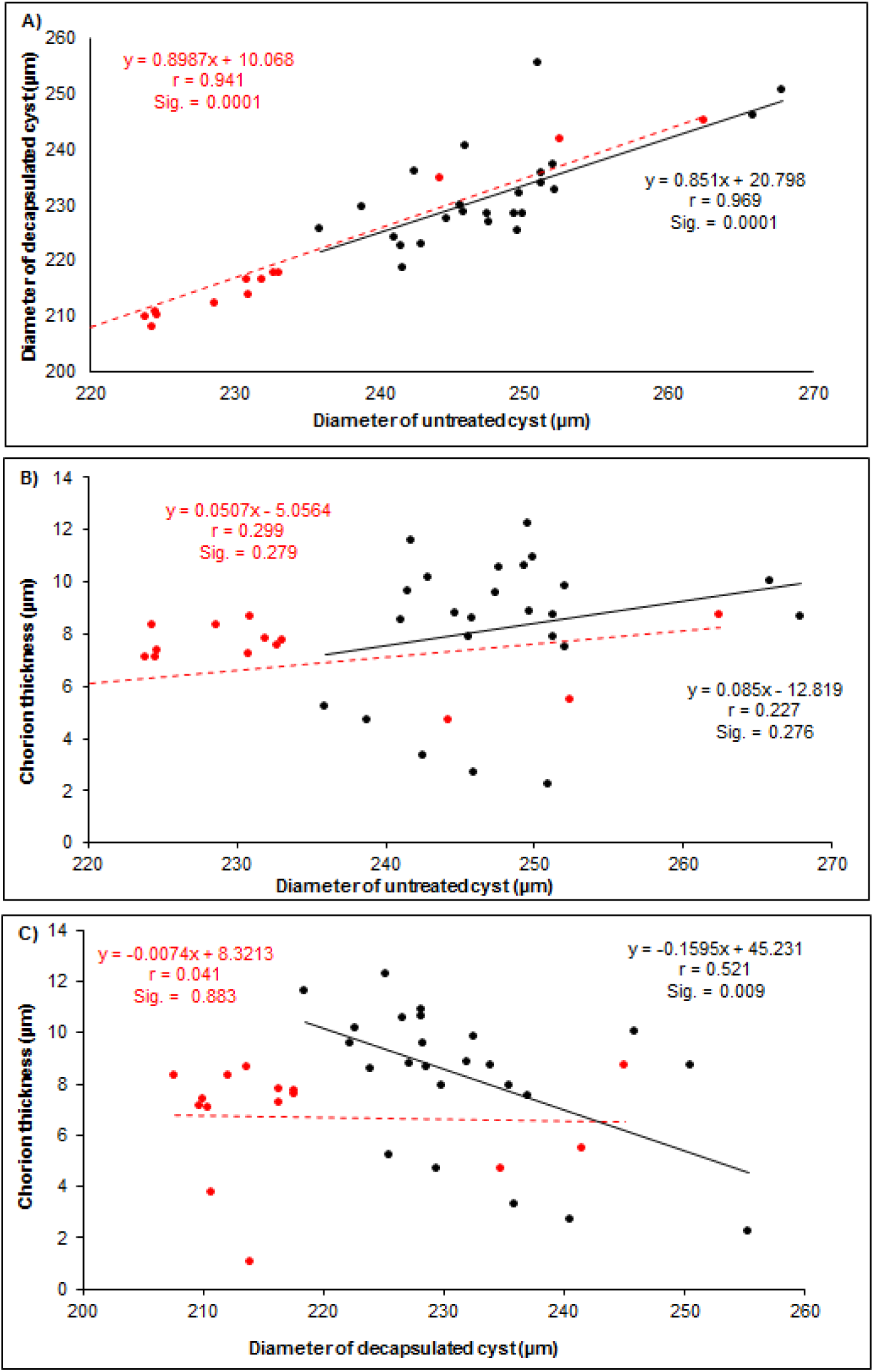
Correlation among biometric characters in native and invasive populations of *A. franciscana* cyst. (Red and black colors have been used for native and invasive populations, respectively). A) Regression between diameters of untreated cyst and decapsulated cyst; B) Regression between diameter of untreated cyst and chorion thickness; C) Regression between diameter of decapsulated cyst and chorion thickness.

## DISCUSSION

The introduction of exotic species to the new habitats can reduce biodiversity and would eventually reorganize the biological communities (Olden et al., 2004; Lodge et al., 2006). Though, almost 1% of the introduced non-natives species become invasive (Williamson, 2006), colonization of exotic species in the new environments have generated extensive economic damage and ecological effects (Pimentel et al., 2005).

Given the economic value of the brine shrimp *Artemia* in fishery and aquaculture, and regarding to the biological potential and high production fitness of American *A. franciscana*, it has been introduced in other continents especially in Asia, including China (Bohai Bay), Vietnam (Vinhchau) and Iran (Nough catchment) (Le et al., 2019; Van Stappen et al., 2020; Eimanifar et al., 2020). Numerous studies have been documented ecological effects of exotic *A. franciscana* which could significantly diminish biodiversity of the native species of *Artemia* (Amat et al., 2007; Scalone & Rabet, 2013; Asem et al., 2018; Eimanifar et al., 2020, Shen et al., 2021). The current survey advances an opportunity, to our knowledge, to perceive cyst size variation of introduced *A. franciscana* in Asia compared to its native populations in the USA.

Generally, *A. franciscana* has represented a wide range of variation in biometry of cyst. Vanhaecke & Sorgeloos (1980) have reported Great Salt Lake (Utah, USA) has a larger cyst diameter compared with San Francisco Bay (California, USA). A study on cyst diameters in the six localities from Colombian Caribbean populations of *A. franciscana* have revealed their cysts characterization were more similar to Great Salt Lake (Camargo et al., 2005). Castro et al. (2006) have reported an extensive cyst variation for Chilean populations (220.5 to 241 μm for untreated cyst and 5.4 to 7.9 μm for chorion thickness) and Mexican populations of *A. franciscana* (200.4 to 292.3 μm for untreated cyst and 2.11 to 10.78 μm for chorion thickness). It has been concluded that intraspecific differentiation in cyst size can be attributed to the seasonal fluctuations in ecological parameters and food availability (Asem et al., 2010).

It has been confirmed that critical environmental conditions (salinity = 300 g/l in 2003) in Urmia Lake could affect cyst characterizations of *Artemia urmiana* to have small size of untreated and decapsulated diameter and thicker chorion layer (Asem et al., 2010). Sankian et al. (2011) have showed that a newly *Artemia* nauplii hatched from wild cysts that have been harvested in optimum ecological condition from Urmia Lake in 1998 (salinity = 180 g/l) had low mortality rate and higher RNA content than cyst sample in 2003 (see Asem et al., 2019). This finding could challenge that whether smaller cysts should have high quality owing to high hatching efficiency. Although, there are multiple reports of cyst size variation of different *Artemia* species/populations, due to the lack of comprehensive studies, our knowledge about interspecific and intraspecific variation of *Artemia* cyst and its relationship with environmental conditions is limited.

Asem & Sun (2014) characterized Chinese parthenogenetic *Artemia* (Crustacea: Anostraca) cysts with different ploidy levels. The results have documented that all three biometrical parameters have positive and significant correlations to each other. By contrast, in the current study there was no significant correlation between diameter untreated cyst and chorion thickness in both native and invasive population. Additionally, invasive populations of *A. franciscana* showed a negative correlation between diameter of decapsulated cyst and chorion thickness, meanwhile parthenogenetic populations have exhibited positive correlation that with increasing diameter of decapsulated cyst, chorion layer was thicker.

With regard to our results, native populations (Great Salt Lake and San Francisca Bay) show dissimilar biometrical patterns. Great Salt Lake batches represent a heterogeneity distribution pattern in PCA, while San Francisca Bay batches are clustered in one group. The reason of this differentiation might be referred to under control conditions in San Francisca Bay saltern(s) as man-made constrictions, so that, Great Salt Lake has been frequently affected via seasonal fluctuations in biotic and abiotic parameters. Although, actual origin of invasive *A. franciscana* populations in non-native environments are unclear (Eimanifar et al., 2014, Asem et al., 2018; Shen et al., 2021), at least it has been confirmed that Vinhchau population in Vietnam have been originated from San Francisco Bay (Le et al., 2019). Even so, Vinhchau population has not been located in the same group with San Francisco Bay which it could be attributed to effect of the environmental conditions. Significant biometrical differentiation exists in the native population of San Francisca Bay (USA).

In conclusion, biometrical characterizations of American *A. franciscana* cyst have exhibited intraspecific variation. Environmental conditions with effect on adaptations to biotic and abiotic parameters could be taken to be an important cause on cyst size variability. Despite of small cyst could not be the only reason for its high quality, the importance of cyst size on hatching efficiency and marketing should be consider. Interspecific and intraspecific variation and regression patterns of *Artemia* cyst necessitate comprehensive studies on biological pathways to understand how bionic and abiotic factors influence cyst size variability.

## ACKNOWLEDGEMENTS

The authors thank Prof. Gilbert Van Stappen (*Artemia* Research Center, Ghent University, Belgium) for preparing *Artemia* cyst samples for this study. This study has been supported by 2021 Hebei Province introduced foreign intelligence project.

